# ZBTB38 requires an extended N-terminal zinc finger network to read mCpG- and discriminate TpG-containing DNA sequences

**DOI:** 10.64898/2026.05.04.722763

**Authors:** Jared Boster, Cooper Gangi, Nicholas O. Hudson, Dallin E. Billings, Brandon Leonel Guerra Castañaza Jenkins, Victoria L. Ding, Bethany A. Buck

**Affiliations:** Department of Chemistry, University of Utah, Salt Lake City, Utah 84112

**Author notes:** Department of Oncological Sciences, Huntsman Cancer Institute, University of Utah School of Medicine, Salt Lake City, UT, USA.

**Keywords:** methyl-CpG binding protein, zinc finger and BTB domain-containing protein, DNA methylation, epigenetics, protein-DNA interaction, transcription

## Abstract

Methylation of cytosine bases in the CpG context (mCpG) is an essential regulatory mechanism cells use to spatially and temporally orchestrate access to genomic regions and mediate transcription. In many diseases, DNA methylation patterns become inappropriately distributed leading to aberrant transcriptional outcomes. Methyl-CpG binding proteins (MBPs) are key epigenetic mediators that selectively recognize mCpG sites, translating these signals into discrete transcriptional responses. ZBTB38 is a zinc finger (ZF) MBP that uniquely harbors two sets of five ZF clusters; each capable of selectively distinguishing mCpG sites. While the cognate DNA sequence and molecular basis for selective mCpG recognition have been defined for the ZBTB38 C-terminal (C-term) ZF domain, the molecular basis for differentiating DNA targets by the N-terminal (N-term) ZF domain remained uncharacterized. Here we report the mCpG-containing consensus sequence for the ZBTB38 N-term ZFs and demonstrate that unlike the other two ZBTB MBP family members ZBTB33 (Kaiso) and ZBTB4, the three shared core ZF domain discriminates against binding to TpG-containing DNA, and that at least one additional N-term ZF is required to stabilize DNA engagement. In addition, we demonstrate that each ZBTB38 ZF domain exhibits preferential target recognition for their respective cognate methylated DNA consensus motif. These findings expand understanding for how ZBTB38 differentially mediates epigenetic-based transcriptional process in normal and disease-state cells by providing new insight into the molecular basis by which the ZBTB38 N-term ZF domain differentiates DNA targets and offering further insight into the interplay between the N- and C-term ZF domains in directing cellular activities.

## Introduction

Controlled access to genomic regions is essential for directing specified cell functions and is in part regulated by epigenetic modifications; reversible chemical changes on the surface of DNA or DNA packaging proteins. One essential epigenetic modification is the addition of a methyl group to the 5-position of cytosine bases in the context of CpG sites (mCpG); required for stabilizing genetic material and regulating gene expression. While this process is central for maintaining normal cellular function, misappropriation of DNA methylation patterns and downstream alterations in the transcriptional landscape have been associated with supporting a number of disease conditions (1-3). This has prompted an interest in defining the molecular mechanisms by which DNA methylation orchestrates chromatin architecture, genomic accessibility and transcriptional outcomes. It is now well recognized that DNA methylation at the level of a single CpG site can deter transcription factor (TF) recognition of consensus binding targets and/or recruit specialized TFs, termed methyl-CpG binding proteins (MBPs), that selectively read mCpGs (4, 5). MBPs subsequently translate mCpG signals into recruitment of protein co-assemblies that alter local chromatin status and transcriptional response (6-13), and as such play a critical intermediary role in regulating epigenetic-driven gene activity.

ZBTB38 (zinc finger (ZF) and BTB (broad complex, tramtrack, bric-á-brac) domain-containing protein 38) represents one of three members of the much larger BTB/POZ (pox ZF) family of TFs that exhibits selective readout of mCpG sites utilizing a conserved set of three ZFs (14) (Fig. 1). The founding ZBTB MBP family member ZBTB33 (Kaiso), as well as ZBTB4 also exhibit bimodal DNA binding, each selectively recognizing mCpG- and TpG-containing DNA consensus sites through their respective shared three ZF domains (14-18) (Fig. 1). Based on the observed binding of ZBTB38 to the *tyrosine hydroxylase* (TH) gene promoter E-box dyad (19), it was also believed to participate in bimodal mCpG and TpG target recognition. Similar to ZBTB33 and ZBTB4, there is mounting evidence that ZBTB38 directly occupies distinct genomic loci (20) and mediates epigenetic-based transcriptional processes associated with normal and disease state functions (21). Indeed, ZBTB38 transcriptional activities have been implicated in normal cell function to mediate proliferation and differentiation (22, 23), apoptosis (24, 25), oxidative stress survival (26), and genomic stability (27). The role of ZBTB38 in modulating disease activity is complex in that its up-regulation has been directly associated with promoting migration/invasion in bladder cancer (28), as well as repressing anti-inflammatory genes that promote rheumatoid arthritis (29). In contrast, re-introduction of ZBTB38 at sites of spinal cord injury redirected autophagy and apoptotic pathways to partially recover damaged tissues (30, 31). Despite its emerging roles in mediating normal and disease-state cellular functions, the molecular intricacies by which ZBTB38 recognizes its methylated DNA targets and translates that epigenetic signal into downstream cellular responses remain to be fully defined.

**Figure 1.**
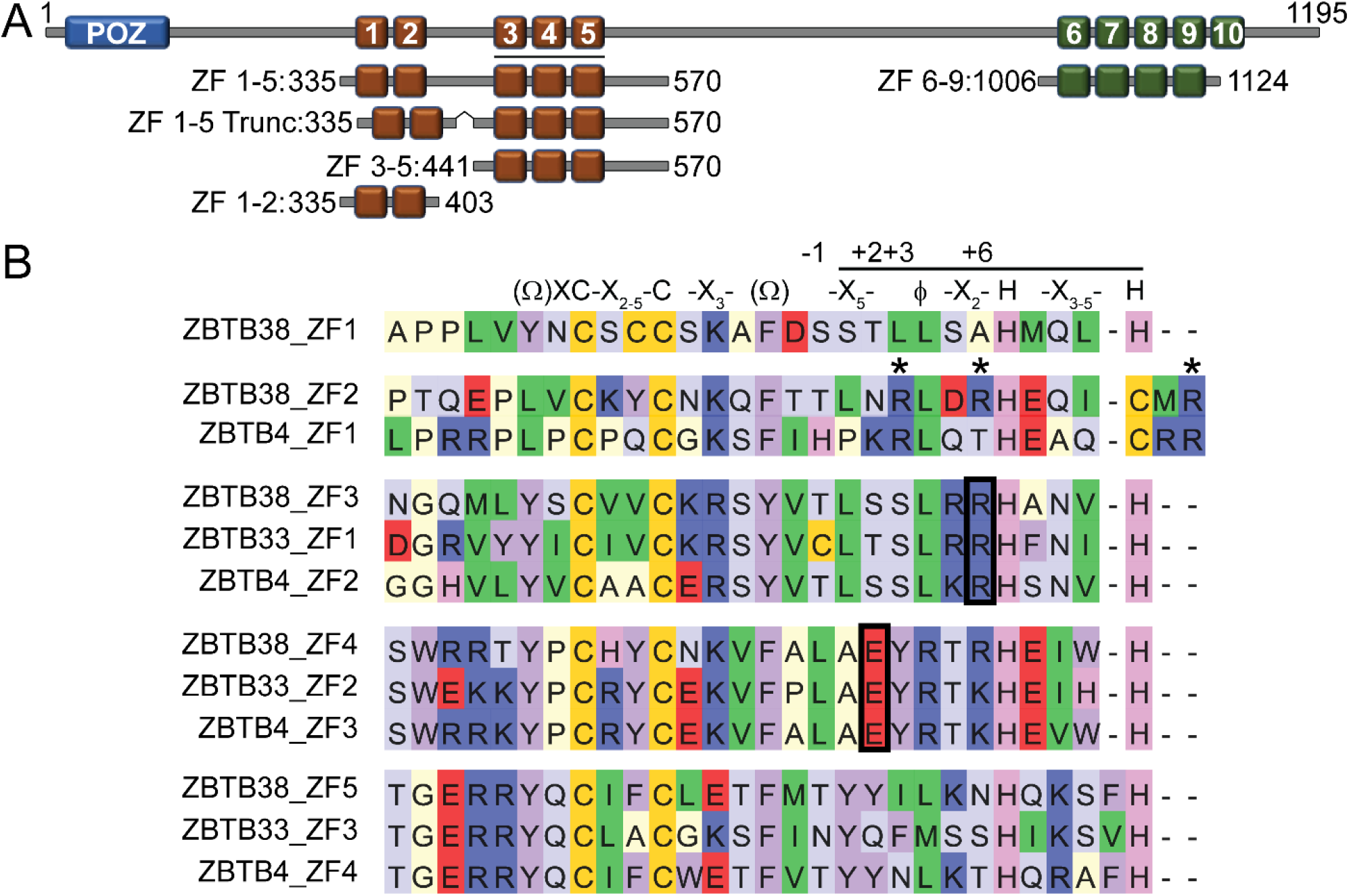
Domain organization of human ZBTB38 and sequence comparison of the N-terminal ZF domains from the three ZBTB MBP family members. *A*, overall domain organization of ZBTB38; the various constructs designed around the N-terminal ZF region (*red boxes*) utilized in this investigation are depicted. The three N-terminal ZFs conserved among the ZBTB MBP family members and responsible for selective methylated DNA read out are underlined. The minimal C-terminal ZF construct (ZFs 6-9; *green boxes*) previously identified to selectively bind methylated DNA utilized in these studies is also depicted. The *blue box* indicates the ZBTB38 BTB/POZ protein-protein interaction domain. *B*, sequence comparison for the N-terminal ZF regions of human ZBTB38 (UniProt ID: Q8NAP3), ZBTB33 (UniProt ID: Q86T24) and ZBTB4 (UniProt ID: QP91Z0). The canonical C_2_H_2_ ZF consensus sequence is shown above the alignment with X, Ω and φ representing any amino acid, F/Y, and a hydrophobic residue, respectively. The helix region is designated with a line and highlights canonical amino acid positions involved in base-specific DNA recognition (-1, +2, +3 and +6). Black boxed residues indicate locations of conserved arginine and glutamate residues known from high-resolution structures to participate in base-specific hydrogen bonding interactions at mCpG sites. In ZBTB38 ZF 2, the arginine residues believed to contribute to DNA binding are indicated by asterisks.

Of the three ZBTB MBP family members, ZBTB38 uniquely harbors two spatially separated five ZF clusters, one localized toward the N-terminal (N-term) BTB/POZ domain, and one toward the C-terminal (C-term) end. With the exception of ZF 2 which is of a C_2_HC type, all ZBTB38 ZFs adopt canonical C_2_H_2_ folds (Fig. 1B), though C^2^HC type ZFs can adopt secondary structural folds similar to C_2_H_2_ ZFs and engage with DNA (32-34). While the ability to selectively distinguish mCpG-containing DNA was originally assigned to the three shared ZF domain (ZFs 3-5) (14, 25, 35), the function(s) of the remaining seven ZFs remained uncharacterized until recently. Specifically, we determined that the ZBTB38 C-term ZF domain also selectively recognizes methylated DNA (termed the methylated C-term ZBTB38 binding site (mCZ38BS)) utilizing a novel molecular mechanism of readout (36, 37). Importantly, cell-based studies indicated that depending on the promoter context, the ZBTB38 N- and C-term ZF domains may work independently or synergistically to modulate transcription (20, 37), though the interplay between these ZF domains in directing cellular activities remains to be fully elucidated. Further, the role(s) of ZBTB38 ZFs 1-2, as well as the molecular basis of DNA readout by the N-term ZFs remained undefined.

Here we utilized an unbiased *in vitro* selection strategy to define the N-term ZF methylated DNA consensus motif (termed the methylated N-term ZBTB38 binding site (mNZ38BS)). Surprisingly, this investigation revealed that ZBTB38 required an extended N-term ZF network to select a cognate methylated DNA target and failed to identify a TpG-containing consensus sequence. Significantly, recent cell-based ZBTB38 ChIP-seq analysis in HeLa-S3 cells corroborated our findings by not only *de novo* identifying two independent methylated DNA consensus motifs that correspond to our respective *in vitro* selected mCZ38BS and mNZ38BS but similarly failing to identify a TpG-containing DNA target (20). Further characterization of the N-term ZBTB38 ZF interactions with DNA demonstrated that ZBTB38 utilizes at least one additional N-term localized ZF outside of the shared core ZF domain to stabilize DNA engagement, though selection of mCpG over CpG and the ability to discriminate TpG-containing DNA sequences is facilitated through the shared three ZF domain (ZFs 3-5). Finally, through comparative binding investigations, we confirmed that each of the ZBTB38 N- and C-term ZF domains exhibit preferential target recognition for their respective cognate methylated DNA consensus motif. Together, these findings expand understanding for how ZBTB38 differentially mediates epigenetic-based transcriptional process by providing new insight into the molecular basis by which the ZBTB38 N-term ZF domain recognizes DNA, assigning function to the additional N-term ZFs, and offering further perspective on the interplay between the ZBTB38 N- and C-term ZF domains in target DNA recognition.

## Results

### ZBTB38 utilizes an extended N-terminal ZF network to selectively distinguish methylated DNA and discriminated TpG-containing DNA sequences

While ZBTB38 shares a conserved set of three N-term localized ZFs (ZFs 3-5) with ZBTB33 and ZBTB4 responsible for highly selective methyl-CpG (mCpG) DNA recognition (Fig. 1), the observation that ZBTB38 occupies preferential genomic targets and orchestrates distinct cellular functions (20, 21, 38) prompted us to question whether the two additional N-term ZFs (ZFs 1-2) may also contribute to DNA binding. Further, upon initiation of these investigations, the high-affinity DNA target(s) for the N-term ZBTB38 ZF domain had yet to be defined. To investigate this, an electrophoretic mobility gel shift assay-systematic expansion of ligands by exponential enrichment (EMSA–SELEX) strategy was employed (39). Based on the bimodal DNA recognition exhibited by both ZBTB33 and ZBTB4 (14-18), as well as prior literature evidence that ZBTB38 may also recognize a TpG-containing DNA sequence (19), we performed the randomized selection in parallel utilizing methylated and non-methylated DNA pools. In addition, N-term ZBTB38 ZF constructs designed around the conserved core three ZFs (ZF 3-5) and encompassing all five ZFs (ZF 1-5) were utilized in parallel with each randomized DNA pool (Fig. 1A). Unexpectedly, and in notable contrast to ZBTB33 and ZBTB4, a consensus motif could only be isolated from the methylated DNA pool and only when all five N-term ZBTB38 ZFs were present (Fig. 2A, Table S1). Importantly, these findings were recently validated by ZBTB38 ChIP-seq studies in HeLa-S3 cells (20), which *de novo* identified two mCpG-containing genomic consensus motifs, one that matches our previously identified consensus motif for the C-term ZF domain ((mCZ38BS); Fig. 2B) (37), and one that aligns with the consensus motif identified here for the five N-term ZFs (mNZ38BS; Fig. 2A). Significantly, the ChIP-seq studies failed to identify a TpG-containing genomic binding target for ZBTB38 (20), reaffirming our EMSA-SELEX observations.

**Figure 2.**
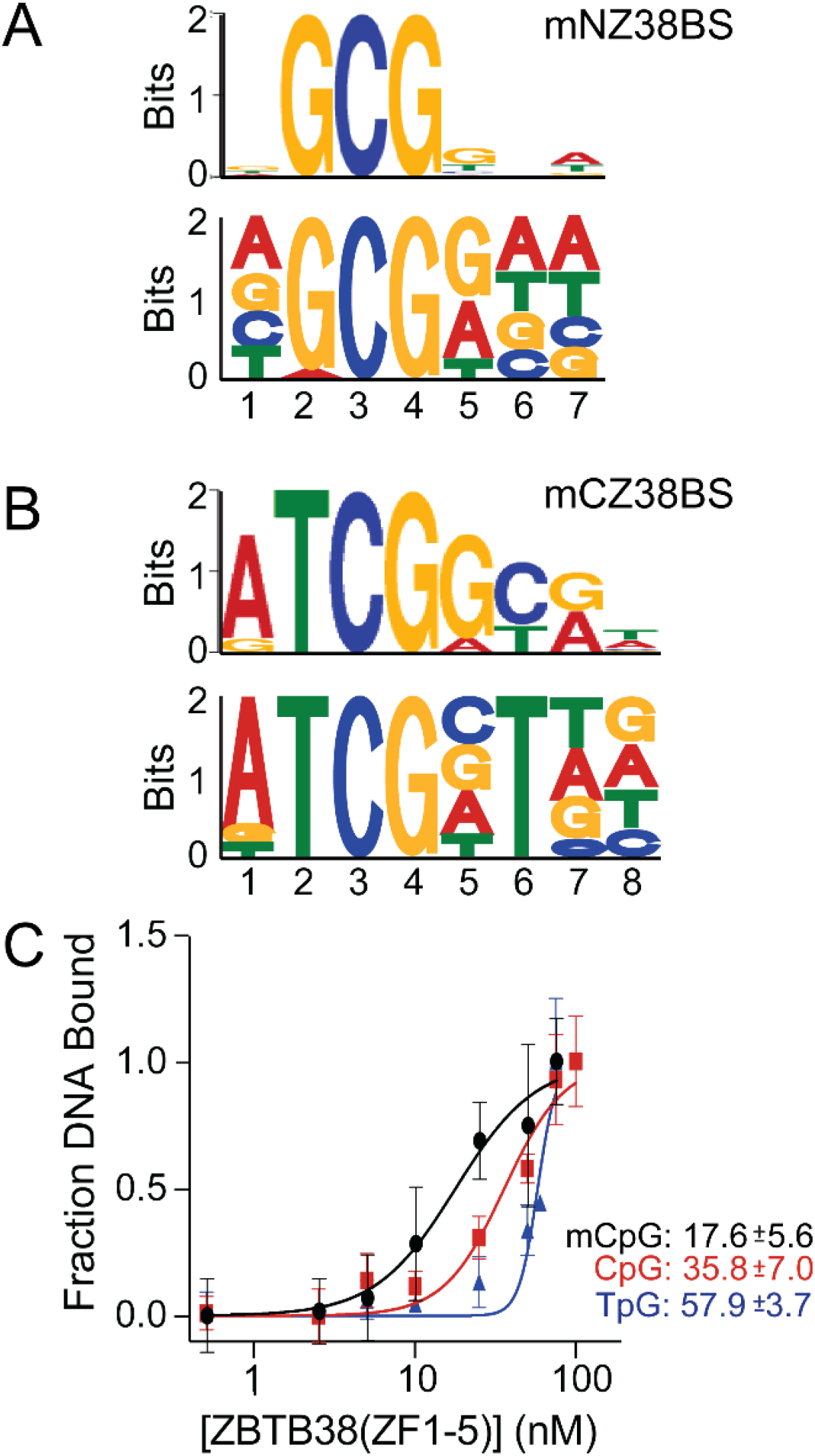
The N- and C-terminal ZBTB38 ZF domains have distinct methylated DNA consensus motifs. *A*, sequence logos for the ZBTB38 N-terminal ZF domain consensus motif derived from EMSA-SELEX (*top*) and by recent ChIP-seq analysis (*bottom*). *B*, sequence logos for the previously determined C-terminal ZF domain consensus motifs derived from EMSA-SELEX (*top*) and by recent ChIP-seq analysis (*bottom*). *C*, comparative binding isotherms for ZBTB38 ZFs 1-5 with mTMPRSS2 (mCpG; *black circles*), TMPRSS2 (CpG; *red squares*), and E-box (TpG; *blue triangles*) DNA sequences.

Our previous investigations identified that the human *TMPRSS2* gene promoter harbored the mCZ38BS and was occupied by the ZBTB38 C-term ZF domain in HEK293 cells in a methyl-dependent manner (37). These previous cell-based studies also suggested that depending on the cellular context, the N- and C-term ZF domains may work independently or synergistically to mediate transcriptional outcomes at a given promoter (37); which has now been reaffirmed on a more global genomic scale (20). Given the importance of the *TMPRSS2* gene promoter in driving a significant proportion of prostate cancer tumors (40-44), we bioinformatically mined this promoter for the mNZ38BS. From this analysis we identified genomic loci matching both the mNZ38BS and mCZ38BS (Fig. S1), the former of which was utilized to design a DNA target (mTMPRSS2; Table S2) to further investigate binding interactions by the ZBTB38 N-term ZF domain.

Quantitative EMSA analysis confirmed that the five N-term ZF domain preferentially recognizes methylated DNA, exhibiting a 2-fold improved binding for mCpG (K_d_ = 17.6 ± 5.6 nM) over CpG (K_d_ = 35.8 ± 7.0 nM) (Fig. 2C, Fig. S2). To comparatively evaluate the ability of this ZF domain to bind a TpG-containing DNA target, we utilized a sequence designed from the previously observed *tyrosine hydroxylase* gene promoter E-box dyad (E-box; Table S2) (19). Consistent with the EMSA-SELEX and ChIP-seq analyses, and in distinct contrast to ZBTB33 and ZBTB4, both of which recognize mCpG and TpG-containing DNA targets with similar affinity (15, 16, 18), ZBTB38 ZFs 1-5 exhibit a 3.3-fold weaker binding for TpG (K_d_ = 57.9 ± 3.7 nM) relative to mCpG, and a 1.6-fold weaker binding than CpG (Fig. 2C, S2). Collectively, these findings suggest that relative to ZBTB33 and ZBTB4, the shared N-term ZBTB38 ZFs uniquely harbor an ability to discriminate TpG-containing DNA targets and require an extended N-term ZF network to achieve optimal DNA binding.

### ZBTB38 ZFs 1-2 participate in non-specific DNA binding while ZFs 3-5 are responsible for selective DNA readout

To differentially compare the ability of the various ZBTB38 N-term ZFs to recognize mTMPRSS2 (Fig. 1A), we utilized a combined solution NMR and EMSA analysis. To evaluate the roles of ZFs 1-2 in DNA binding, a construct encompassing only these ZFs (residues 335-403; ZF 1-2) was designed (Fig. 1A). ^1^H-^15^N HSQC analysis of ZFs 1-2 in complex with mTMPRSS2 resulted in a well dispersed spectrum exhibiting chemical shift perturbations, indicative of a direct binding interaction (Fig. 3A). In addition, ZF 1-2 complexation of mTMPRSS2 resulted in the appearance of three arginine guanidinium side chain resonances that only appear under neutral pH conditions when these protons are stabilized in non-bonding interactions (e.g. nucleobase hydrogen bonding or phosphate backbone electrostatic interactions) with DNA (Fig 3A/3C). Consistently, sequence analysis of ZFs 1-2 indicated the presence of three arginine residues, all of which are localized in ZF 2 (Fig. 1B), suggesting that at least this ZF participates in DNA binding interactions. However, the minimal observed chemical shift perturbations for the base imino resonances of mTMPRSS2 as evaluated by 1D ^1^H NMR analysis (Fig. 3B) indicated that ZBTB38 ZFs 1-2 participate in non-specific DNA binding interactions.

**Figure 3.**
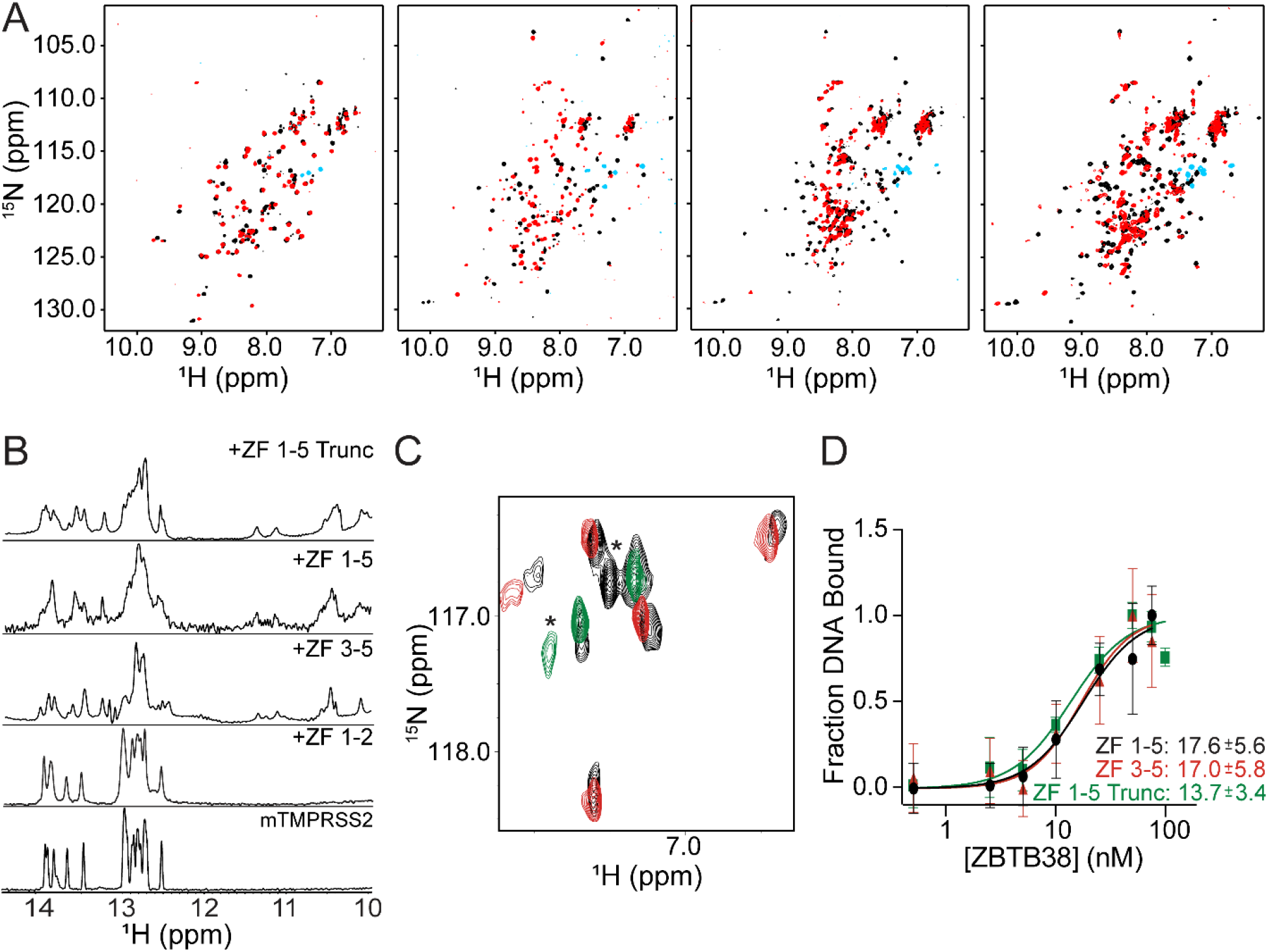
ZBTB38 utilizes more than the three conserved N-terminal ZFs for DNA binding. *A*, ^1^H-^15^N HSQC spectra of the various ZBTB38 N-terminal ZF constructs free (*black spectra*) and in complex with mTMPRSS2 (*red spectra*). *Blue* cross peaks denote aliased arginine guanidinium side-chain resonances that appear upon mTMPRSS2 binding at neutral pH. *B*, 1D ^1^H NMR spectra of the mTMPRSS2 imino proton region free and in complex with the various ZBTB38 N-terminal ZF constructs. *C*, zoom in and overlay of the ^1^H-^15^N HSQC aliased arginine guanidinium side-chain resonance spectral region for ZFs 1-5 (*black*), ZFs 1-2 (*green*), and ZFs 3-5. Asterisks denote the chemical shift change for what is believed to be the terminal arginine residue (Arg-395) in ZF 2, which would experience a different local chemical environment between the ZF 1-2 and ZF 1-5 protein constructs. *D*, comparative binding isotherms for ZF 1-5 (*black circles*; replotted from Fig. 2C for comparison), ZF 3-5 (*red triangles*), and ZF 1-5 truncate (*green squares*) with mTMPRSS2 DNA and associated *K*_*d*_ values.

Similarly, to independently evaluate the role of the three conserved N-term ZFs, a construct encompassing only these ZFs (residues 441-570; ZF 3-5) was designed (Fig. 1A). Consistent with this ZF domain harboring the core arginine and glutamate residues required for base-specific readout of the mCpG palindrome (Fig. 1B) (5), NMR analysis of ZFs 3-5 in complex with mTMPRSS2 yielded spectra with a significant increase in chemical shift perturbations for all three ZFs, as well as the mTMPRSS2 base iminos as evidenced by the ^1^H-^15^N HSQC and 1D ^1^H spectra, respectively (Fig. 3A/3B). In preparation to comparatively evaluate DNA binding by all five N-term ZBTB38 ZFs (335-570; ZF 1-5), it was observed that this construct had a significant decrease in solubility and required the addition of detergent to maintain it in solution at the concentrations required for NMR spectral analysis. ^1^H-^15^N HSQC analysis of all five N-term ZFs (ZF 1-5) in complex with mTMPRSS2 resulted in a nearly complete loss of resonances localized in ordered structured regions of the protein and a line broadening increase for mTMPRSS2 imino resonances in the 1D ^1^H spectrum (Fig. 3A/3B). Nevertheless, the observed chemical shift perturbations for mTMPRSS2 imino resonances (Fig. 3B), as well as the appearance of arginine guanidinium side chain resonances belonging to both ZF 1-2 and ZF 3-5 constructs (Fig. 3C), indicated that ZBTB38 utilized an extended N-term ZF region to engage DNA.

Further evaluation of the free ZF 1-5 ^1^H-^15^N HSQC spectrum showed a significant number of resonances within the intrinsically disordered region (IDR) of the spectra belonging to the ∼40 amino acid linker separating ZFs 1-2 and 3-5, that did not appear to experience chemical shift perturbations or significant line broadening upon addition of mTMPRSS2 (Fig. 3A). This observation suggested that this region was not involved in DNA binding. Thus, a truncated variant excising much of this IDR was designed to fuse the ZF 1-2 and ZF 3-5 constructs together (335-403,441-570; termed ZF 1-5 Trunc; Fig. 1A) and evaluated for DNA binding. Consistent with our observations, removal of the IDR not only improved solubility of the five ZF N-term construct, but in complex with mTMPRSS2 yielded a ^1^H-^15^N HSQC spectrum with more uniform cross-peaks and additional chemical shift perturbations relative to ZFs 3-5 (Fig. 3A), indicative of the formation of a more stabilized complex. Quantitative EMSA analysis comparing the binding of mTMPRSS2 by ZF 3-5 (17.0 ± 5.8 nM), ZF 1-5 (17.6 ± 5.6 nM) and ZF1-5 Trunc (13.7 ± 3.4 nM) demonstrated the binding to be similar within error (Fig. 3D, Fig. S3). Taken together these findings definitively affirm that ZBTB38 utilizes an extended N-term ZF network to recognize DNA, with ZFs 1-2 likely contributing stabilizing interactions, while ZFs 3-5 participate in base-specific readout.

### The ZBTB38 N- and C-term ZF domains exhibit preferential binding for their respective consensus motif

While distinct methylated DNA targets have been identified for the ZBTB38 N- and C-term ZF domains, a comparative binding analysis evaluating the ability of each domain to differentiate its own cognate recognition motif over the other has not yet been performed. Thus, we utilized EMSA analysis to quantify the ability of the N- (ZF 1-5) and C-term (ZF 6-9) ZBTB38 ZF domains to recognize their own consensus sequence and that of the other ZF domain. In both cases, each ZF domain exhibited preferential binding for their respective consensus motifs (Fig. 4, Fig. S4). Notably, the measured *K*_*d*_ value for the ZBTB38 C-term domain binding to the mCZ38BS was higher than our previous report (36), though different assay buffer conditions were utilized in this current investigation to allow direct comparison of DNA binding with the N-term ZF domain. Significantly, the N-term ZF domain only differentiated ∼2.3-fold between the mTMPRSS2 (*K*_*d*_ = 17.6 ± 5.6 nM) and mCZ38BS (*K*_*d*_ = 40.6 ± 8.0 nM), while the C-term ZF domain demonstrated a much higher ability to differentiate between the two exhibiting an ∼4-fold difference between mCZ38BS (*K*_*d*_ = 13.8 ± 2.8 nM) and mTMPRSS2 (*K*_*d*_ = 56.2 ± 2.8 nM) recognition. This finding is consistent with the C-term ZF domain requiring specific sequence context outside of the core mCpG palindrome at positions 2 and 6 (Fig. 2B), while the N-term ZF domain only requires a minimal mCpGpC core (Fig. 2A).

**Figure 4.**
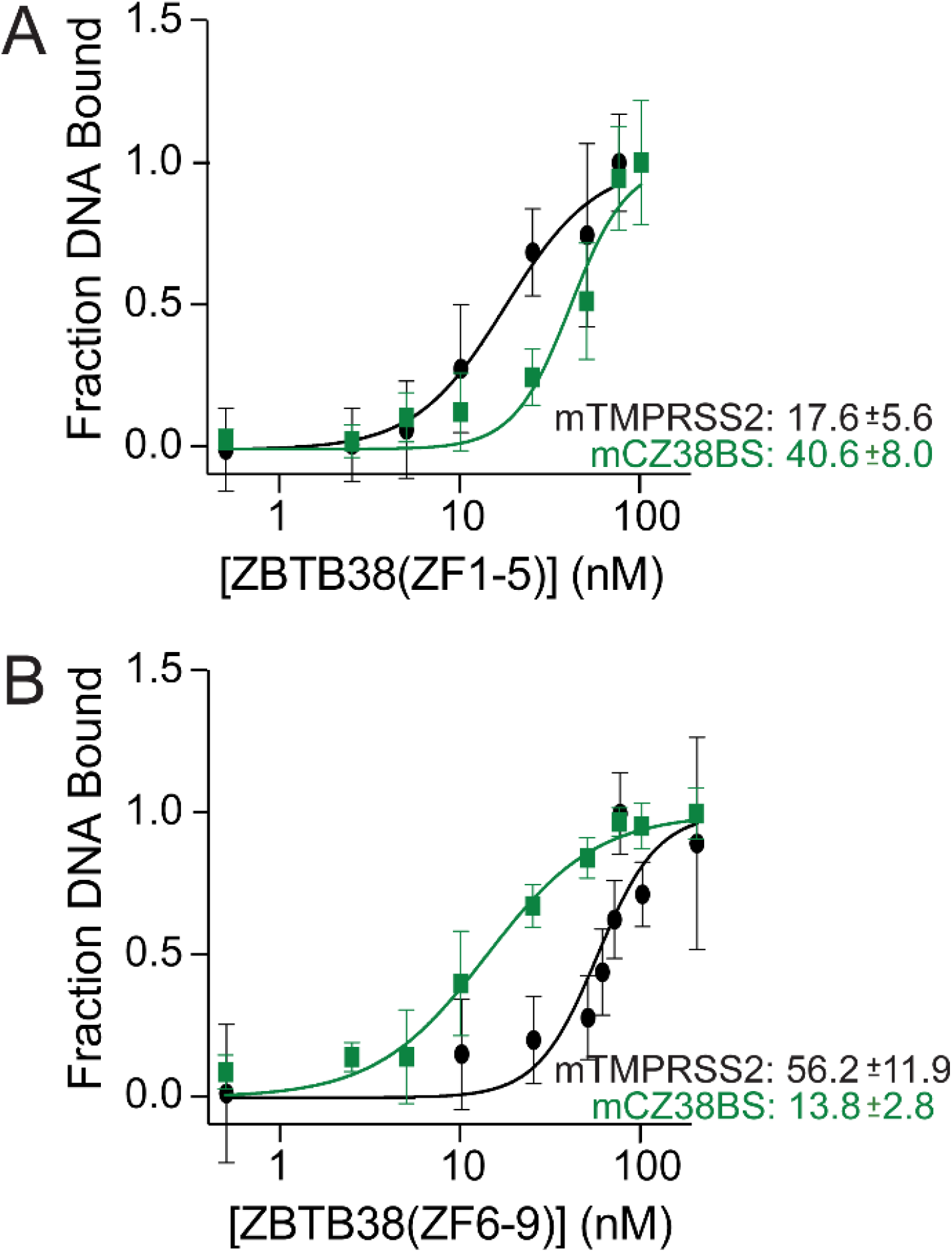
The ZBTB38 N- and C-terminal ZF domains are selective for their respective methylated DNA consensus motif. *A*, comparative binding isotherms for ZBTB38 ZFs 1-5 with mTMPRSS2 (*black circles*; replotted from Fig. 2C for comparison) and mCZ38BS (*green squares*). *B*, comparative binding isotherms for ZBTB38 ZFs 6-9 with mCZ38BS (*green squares*) and mTMPRSS2 (*black circles*).

## Discussion

Here we utilized an unbiased *in vitro* selection strategy to identify the N-term ZBTB38 ZF domain cognate methylated DNA target, termed the mNZ38BS (Fig. 2A). Significantly, the mNZ38BS and our previously unbiased *in vitro* selected mCZ38BS (37) were corroborated by global genomic occupation ChIP-seq experiments in HeLa-S3 cells (20); indicating that our strategy is robustly capable of identifying the physiologically relevant DNA consensus motif for a given protein. This is notable in that our selection strategy also afforded us the ability to determine that unlike ZBTB33 and ZBTB4, the ZBTB38 N-term ZF domain requires an extended ZF network to achieve optimal DNA binding; information that would not be discernable through ChIP-seq. NMR spectroscopic analysis further confirmed the involvement of ZBTB38 ZF 2 in direct DNA binding as evidenced by ^1^H-^15^N HSQC chemical shift perturbations and the appearance of three arginine guanidinium side chains (Fig. 3A/3D). This does not, however, rule out a role for ZBTB38 ZF 1 in also contributing to overall stabilization of DNA binding. NMR analyses indicated that collectively ZFs 1-2 participate in non-specific DNA binding interactions (Fig. 3A/3B), so it may be that ZF1 contributes non-bonding interactions with the DNA backbone, or alternatively, ZF 1 may directly interact with the other N-term ZFs, indirectly stabilizing the DNA binding interaction. In addition, while all three ZBTB MBP family members share a set of three ZFs responsible for base-selective readout of mCpG sites (Fig. 1B), in contrast to ZBTB33 and ZBTB4, this domain in ZBTB38 has the unique ability to discriminate against TpG DNA recognition. High-resolution structural determination for the ZBTB38 N-term ZF domain in complex with its identified cognate methylated DNA target will provide necessary insight for defining the role(s) of ZFs 1-2 in enhancing DNA binding, as well as illuminate the molecular basis by which ZBTB38 uniquely discriminates against TpG-containing DNA binding.

Of all the MBPs identified and characterized to date, ZBTB38 remains the only one to have two spatially separated ZF domains that each selectively recognize distinct methylated DNA sequences. Our findings reaffirmed that each ZBTB38 ZF domain exhibits preferential binding for its own cognate mCpG-containing site, though the C-term ZFs 6-9 do so with a higher selectivity relative to the N-term ZFs 1-5 (Fig. 4). This is unsurprising given that the C-term ZFs exhibit more extensive base-specific sequence readout beyond the core mCpG palindrome. Similar to ZBTB38 ZFs 3 and ZF 4, ZFs 7 and 8 each contribute the requisite amino acids now known to facilitate ZF recognition of core mCpG bases (5), Lys-1055 (makes a water-mediate hydrogen bond with a mCpG guanine) and Glu-1079 (makes classical and CH…O hydrogen bonds with a mC), respectively (36). Outside of the core mCpG palindrome, Asn-1052 (ZF 7) makes a bifurcated hydrogen bond interaction with the adenosine paired with the thymine localized at position 2 in the consensus motif (Fig. 2B); a key component in selective targeting of this DNA sequence. Indeed, as demonstrated here (Fig. 4B) as well as our prior investigation (36), substitution of position 2 from a T:A step (mCZ38BS) to a C:G step (mTMPRSS2) resulted in a loss in DNA binding. Also notable, our previous EMSA-SELEX investigations using a randomized methylated DNA pool selected a motif requiring a methylated pyrimidine (either mC or T) to occupy position 6 of the consensus sequence (37), however, the recent ZBTB38 ChIP-seq studies determined that in the genomic context this position is exclusively a thymine (20) (Fig. 2B). Here, consistent with our previous findings (37), substitution of a methylated pyrimidine (mCZ38BS) for a cytosine (mTMPRSS2) in position 6 contributed to a loss in DNA binding capability (Fig. 4B).

In contrast, the N-term ZFs recognize a much smaller consensus motif mCpGpC. Based on the high-resolution structure of ZBTB33 in complex with its methylated DNA target (16), it is expected that the conserved Arg-474 (ZF 3) and Glu-501 (ZF 4) in ZBTB38 make respective base-specific classical and CH…O hydrogen bonding interactions with the core mCpG guanine and methyl-cytosine. The requirement for the additional 3′-C:G step could be due to Arg-474 making cross-strand hydrogen bonds with the mCpG core guanine and the paired guanine in the 3′-C:G step, as previously observed for the ZBTB33:methylated DNA complex (16). Here, substitution of the C:G step (mTMPRSS2) to a G:C step (mCZ38BS) only moderately decreased DNA binding (Fig. 4A). Together, these findings suggest that while each ZBTB38 ZF domain prefers its own methylated DNA consensus, in the larger genomic context, the N-term ZF domain has more versatility to directly occupy a broader target recognition landscape. Indeed, the ChIP-seq investigations identified a larger number of ZBTB38 genomic occupation sites harboring the N-term ZF domain consensus sequence relative to that of the C-term ZFs (20).

Nevertheless, the full biological implications of the ability of ZBTB38 to independently and simultaneously readout two distinct methylated DNA sequences remains to be fully realized. However, it is notable that in the broader cellular context, ZBTB38 genomic occupation loci can harbor either the N- or C-term consensus motif, or both (20), further expanding the concept that these domains may work independently or synergistically to mediate transcription (37). In addition, ZBTB38 exists as a homodimer through its BTB/POZ domain, which in conjunction with its modularity in methylated DNA readout may provide the capability to orchestrate long-range chromatin organization to direct transcriptional outcomes. Indeed, ZBTB38 was shown by ChIP-seq to have enriched genomic occupation at enhancer elements (20), and has been identified as a candidate master TF that functions at super enhancer regions (45). Nevertheless, revealing the intricacies by which the two ZBTB38 ZF domains orchestrate transcriptional processes will require further cell-based investigations.

### Experimental procedures

#### Sample Preparation

An initial N-term human ZBTB38 ZF domain construct encompassing resides (335-586; UniProt ID: Q8NAP3) was PCR amplified from a human brain cDNA library (Life Technologies - Thermo Fisher Scientific) and cloned into the pET-21d *E. coli* overexpression vector (Novagen). All other continuous ZBTB38 N-term ZF-containing constructs (Fig. 1A) were PCR amplified using ZBTB38(335-586) as a template and cloned into the pET-21d *E. coli* overexpression vector. The N-term five ZF ZBTB38 construct deleting amino acids 404-440 within the IDR separating ZFs 1-2 and 3-5 (ZF 1-5 Trunc, Fig. 1A) was generated using a one-step PCR fragment deletion strategy (46). Specifically, the truncate variant was generated by PCR with Taq Polymerase (Life Technologies - Thermo Fisher Scientific), using pET-21d-ZBTB38(335-570) as a template with forward (5′-CCATTCCTGGAGGAAGTACTTTGCCAGACACGGACC-3′) and reverse (5′-AGTACTTCCTCCAGGAATGGGCATGTGGCTTGAC-3′) primers: 1 × 95 °C for 2 min;10 × 95 °C for 30 s, 55 °C for 30 s, 68 °C for 30 s, and 68 °C for 1 min; and 1 × 68 °C for 7 min. All primers for the above described PCR reactions were synthesized by the University of Utah DNA/Peptide Synthesis Core Facility and used without further purification.

^15^N-labeled protein was overexpressed in BL21-CodonPlus (DE3)-RIL *E*. coli cells (Agilent) in M9 minimal medium supplemented with 1.0 g ^15^NH_4_Cl/1L of culture (Cambridge Isotope Labs) and 150 μM ZnSO_4_/1L of culture during IPTG induction (1 mM) at 16 °C for 16 h. After cell culturing, pelleted cells were resuspended in lysis buffer (20 mM Tris (pH 8.0), 8 M urea, 200 mM Arg–HCl and 10 mM DTT), incubated on ice for 10 min, lysed by sonication, and centrifuged. Pooled lysate supernatant was loaded onto a pre-equilibrated (A: 20 mM Tris (pH 8.0), 8 M urea, 200 mM Arg–HCl and 2 mM DTT) 100 ml SP Sepharose FF column (Cytiva) using a peristaltic pump and eluted at 3 ml/min using the following gradient to 1 M NaCl (B: 20 mM Tris (pH 8.0), 8 M urea, 200 mM Arg–HCl and 2 mM DTT, 1 M NaCl): 98% A/2% B at 0.2%/min; 90% A/10% B at 0.5%/min; hold at 90% A/10% B for 10 min; 80% A/20% B at 0.5%/min; 30% A/70% B at 0.8%/min; 0% A/100% B at 1.2%/min. Protein fractions were pooled and diluted 3x in H2O/0.01% TFA and purified by RP-HPLC on a 55-g, 15-μm, 300-Å, C18 PuriFlash column (Interchim) pre-equilibrated with 90% A (H2O/0.1% TFA)/10% B (ACN/0.1% TFA). Bound protein was eluted with a variable gradient (2%/min to 80% A/20% B; 1%/min to 70% A/30% B; 0.3%/min to 60% A/40% B; 1%/min to 50% A/50% B) and lyophilized for 5–6 days. Protein purity was verified by SDS–PAGE. The ZBTB38 C-term ZF domain construct used in these studies (ZFs 6-9) was cloned, overexpressed in *E*. coli, and purified as described previously (36).

All ZBTB38 ZF domain constructs were refolded by dissolving lyophilized protein in 1.2 ml denaturing buffer (10 mM Tris (pH 7.0), 8 M urea, 10 mM DTT) and rapidly diluting to 0.5 M urea by adding denatured protein solution into 20 ml refolding buffer (10 mM Tris (pH7.0), 10 mM DTT) containing 1.1 molar equivalents ZnSO_4_ per zinc site while stirring. Refolded protein was concentrated in an Amicon stirred cell (MilliporeSigma) to 500 µL or 1 mL and exchanged into NMR buffer (10 mM Tris (pH 6.8), 1 mM tris(2-carboxyethyl)phosphine (TCEP, Gold BioTechnology, Inc.), 0.005% NaN_3_, 10% D_2_O (for ZFs 1-2, ZFs 3-5, ZFs 1-5 Trunc, ZFs 6-9), or 10 mM Tris (pH 6.8), 1 mM TCEP, 0.005% NaN_3_, 75 µM n-dodcyl-β-D-maltoside (DDM, AdipoGen Life Sciences), and 10% D_2_O (for ZFs 1-5) utilizing a Nap-5 or Nap-10 Sephadex G25 column (Cytiva).

All oligonucleotides utilized for EMSA and solution NMR spectroscopy were synthesized by the University of Utah DNA/Peptide Synthesis Core facility and are reported in Table S2. DNA oligonucleotides were further purified by ion pair reversed phase HPLC at 70 °C using an XBridge Premier Oligo BEH C18 column (Waters) starting at 98% A (100 mM triethylammonium acetate (TEAA, pH 7.0) in ultrapure water/2% B (100 mM TEAA (pH 7.0) in acetonitrile) at 0.5 ml/min with the following gradient: 92% A/8% B at 3%/min; 88% A/12% B at 0.8%/min; 76% A/24% B at 0.5%/min; 50% A/50% B at 4%/min; 20% A/80% B at 12%/min; 1% A/99% B at 4%/min; hold at 1% A/99% B for 10 min; 98% A/2% B 9.8%/min. Peaks were monitored at 260 and 300 nm. Eluted HPLC fractions were dialyzed against ultrapure water using regenerated cellulose 3500 MWCO dialysis tubing (Fisher Scientific), lyophilized, resuspended in 10 mM Tris and pH adjusted to 7.0. Duplex DNA was formed by mixing complementary oligonucleotide strands in stoichiometric amounts, heating to 90 °C, and annealing by slow cooling to room temperature.

#### EMSA-SELEX

The SELEX DNA template and primers were synthesized by the University of Utah DNA/Peptide Synthesis Core Facility and used without further purification. The SELEX template was a 66-mer oligonucleotide containing a centralized 19-mer randomized sequence (CTAGCCATGGGTGAAGTCAGTTAT(N^19^)ACTCACCCTCCAGAAGCTTGGAC). The randomized duplex pool was generated by PCR with Taq Polymerase (Life Technologies - Thermo Fisher Scientific), a 5′-Cy3-labeled forward primer (SELEX-F: Cy3-CTAGCCATGGGTGAAGTCAGTT) and reverse primer (SELEX-R: GTCCAAGCTTCTGGAGGGTGAG): 1 × 95 °C for 2 min; 4 × 95 °C for 30 s, 60 °C for 15 s and 68 °C for 15 s; and 1 × 68 °C for 1 min. The randomized duplex pool was gel purified and concentrated using a DNA Clean and Concentrator-5 column (Zymo). For non-methylated DNA SELEX, the purified randomized duplex pool was used directly. For methylated DNA SELEX, the purified randomized duplex pool was methylated using M.SssI and S-adenosylmethionine (Life Technologies - Thermo Fisher Scientific) and purified using a DNA Clean and Concentrator-5 column (Zymo) prior to use in each round of SELEX. For each SELEX round, 4 nM of either nonmethylated or methylated randomized DNA pool was incubated with 80, 40, or 20 nM of ZBTB38(335-586; ZFs 1-5) or ZBTB38(441-570; ZFs 3-5) depending on the selection round (decreased in later rounds to apply selection pressure) in binding buffer (10 mM Tris (pH 7.0), 1 mM TCEP, 0.005% NaN_3_, 150 mM NaCl, 100 μg/ml BSA and 10% (w/v) sucrose) at room temperature for 30 min. Protein-bound DNA was separated by 8% (w/v) non-denaturing PAGE and imaged on a Typhoon Biomolecular Imager (Cytiva) at 425 nm. Super-shifted DNA bands were excised, gel extracted and purified using a DNA Clean and Concentrator-5 column (Zymo). Isolated and purified selection pools were PCR amplified: 1 × 95 °C for 2 min;10 × 95 °C for 30 s, 60 °C for 15 s and 68 °C for 15 s; and 1 × 68 °C for 1 min. After 8, 10 or 14 rounds of selection, 50 ng of each final selected DNA pool was PCR amplified using a non-Cy3-labeled SELEX-F primer. PCR-amplified SELEX products were ligated into a TOPO-TA Cloning vector (Life Technologies - Thermo Fisher Scientific) and transformed into One Shot TOP10 chemically competent cells (Life Technologies - Thermo Fisher Scientific). Plasmids were purified by QIAprep Spin Miniprep Kit (QIAgen) and sequenced (University of Utah DNA Sequencing Core Facility). Motif analysis was performed using the MEME suite (47) (Table S1). Identification of mNZ38BS and mCZ38BS loci within the characterized TMPRSS2 promoter (48) (Fig. S1) were identified by utilizing a weight matrix search (ScoreSequence script within the USeq package) (49) derived from the EMSA–SELEX MEME motif analysis.

#### Solution NMR spectroscopy

N-term ZBTB38 ZF:mTMPRSS2 complexes (100 µM) were made by first diluting the mTMPRSS2 in 7 mL of NMR buffer (10 mM tris (pH 6.8), 1 mM TCEP, 0.005% NaN_3_, 10% D_2_O (ZF 1-2, ZF 3-5, and ZF 1-5 Trunc) or 10 mM Tris (pH 6.8), 1 mM TCEP, 0.005% NaN_3_, 75 µM DDM, and 10% D_2_O (ZFs 1-5), and slowly adding protein dropwise while stirring. In order to prevent higher order aggregation, all complexes were formed using a 0.9:1 N-term ZBTB38 ZF:mTMPRSS2 concentration ratio. Complex samples were concentrated to 350 µL in an Amicon stir cell (MilliporeSigma) under argon gas, centrifuged and transferred to a Shigemi NMR tube. All NMR spectra were recorded at 298 K on a Varian VNMRS 800-MHz spectrometer. All data were processed using NMRPipe (50) and analyzed with NMRView (51) within the NMRbox suite (52).

#### EMSA

To determine *K*_*d*_ values, 2 nM of duplex DNA (Table S2) were incubated with increasing concentrations of various ZBTB38 ZF domain constructs in 20 µL binding buffer (10 mM Tris (pH 7.0), 150 mM NaCl, 1 mM TCEP, 0.005% NaN_3_, 25 µg/mL bovine gamma globulin (BGG, Gold Biotechnology, Inc.), 2.5 µM DDM, and 10% (w/v) sucrose). Reaction mixtures were incubated at room temperature for 10 min, then separated by electrophoresis in 8% 1x tris borate (pH 8.3) nondenaturing polyacrylamide gels at 100 V for 45 min. The gels were soaked in 1x tris borate supplemented with 1x SYBR Gold Nucleic Acid Gel Stain (Life Technologies - Thermo Fisher Scientific) and visualized on an Amersham Biosciences Typhoon Biomolecular Imager (Cytiva) at 495 nm. Gel band intensities were quantified by ImageQuant™ TL software (Version 10.1, Cytiva, Uppsala, Sweden). Notably, at higher concentration ratios of the various ZBTB38 N-term ZF domains relative to DNA, the system would form higher order solubilized aggregates in solution that failed to enter the gels. Only lanes where sufficient sample entered the gel were included in the analysis. Gel images are included in Fig. S2-S4. *K*_*d*_ values were obtained by fitting averaged triplicate fraction-bound DNA data as a function of protein concentration to a Langmuir isotherm (GraphPad Prism v10.0), where the maximal fraction bound value was set to 1.

## Supporting information

This manuscript contains Tables S1-S2 and Figs. S1-S4.

## Acknowledgements

We are thankful to Dr. P. F. Flynn for many helpful discussions, and to Dr. P. Oblad, D. Edwards, and N. Wojnowski for their support in the David M. Grant NMR Center. All NMR results were recorded in the David M. Grant NMR Center, a University of Utah core facility. DNA oligonucleotides were synthesized by the DNA/Peptide Synthesis Core Facility, part of the Health Sciences Center Cores at the University of Utah. DNA sequencing was performed at the DNA Sequencing Core Facility, University of Utah.

## Funding and additional information

This work was supported by MCB-1715370 from the National Science Foundation and the University of Utah. National Science Foundation Research Experiences for Undergraduates supported the training and research contributions of D.E.B. (NSF REU 2150526). This work was supported by UROP from the Office of Undergraduate Research at the University of Utah awarded to V.L.D. Funds for construction of the David M. Grant NMR Center and the helium recovery system were obtained from the University of Utah and National Institutes of Health (NIH) awards 1C06RR017539-01A1 and 3R01GM063540-17W1, respectively. NMR instruments were purchased with support of the University of Utah and NIH award 1S10OD25241-01. This study made use of NMRbox: National Center for Biomolecular NMR Data Processing and Analysis, a Biomedical Technology Research Resource (BTRR), which is supported by NIH grant P41GM111135 (NIGMS). The contents of this publication are solely the responsibility of the authors and do not necessarily represent the official views of the NIH.

