## Supplementary material for "ZBTB38 requires an extended N-terminal zinc finger network to read mCpG- and discriminate TpG-containing DNA sequences": SI

Contains supplemental Tables S1-S2 and supplemental Figures S1-S4

**Table S1.** EMSA-SELEX sequencing results for the five N-terminal ZBTB38 zinc fingers with the methylated DNA pool*

| | |
|---|---|
| mNZ38_1 | GTCCAAGCTTCTGGAGGGTGAGTCAAAGGTAAGCTAAT**TGCGATA**ACTGACTTCACCCATGGCTAG |
| mNZ38_2 | GTCCAAGCTTCTGGAGGGTGAGTGGCGTACTCA**TGACGCA**TCATAACTGACTTCACCCATGGCTAG |
| mNZ38_3 | GTCCAAGCTTCTGGAGGGTGAGTATCCCTCCTCC**ACCCGCC**GATAACTGACTTCACCCATGGCTAG |
| mNZ38_4 | GTCCAAGCTTCTGGAGGGTGAGTGGATAATTC**TTCCGCA**CGAATAACTGACTTCACCCATGGCTAG |
| mNZ38_5 | GTCCAAGCTTCTGGAGGGTGAGTGGCACCACT**ATCCGCA**GCTATAACTGACTTCACCCATGGCTAG |
| mNZ38_6 | GTCCAAGCTTCTGGAGGGTGAGTAAGAGCGAGGTT**TCACGCC**ATAACTGACTTCACCCATGGCTAG |
| mNZ38_7 | GTCCAAGCTTCTGGAGGGTGAGTGGACCACAC**AGCGTCA**TCTATAACTGACTTCACCCATGGCTAG |
| mNZ38_8 | GTCCAAGCTTCTGGAGGGTGAGTTCCGGGCACT**GGCGGCA**TAATAACTGACTTCACCCATGGCTAG |
| mNZ38_9 | GTCCAAGCTTCTGGAGGGTGAGTACTCCATTAA**TCCCGCA**ACATAACTGACTTCACCCATGGCTAG |
| mNZ38_10 | GTCCAAGCTTCTGGAGGGTGAGTATCATGTCCCA**TAGCGCC**AATAACTGACTTCACCCATGGCTAG |
| mNZ38_11 | GTCCAAGCTTCTGGAGGGTGAGTGCAACG**CGCGGTT**CCATACATAACTGACTTCACCCATGGCTAG |
| mNZ38_12 | GTCCAAGCTTCTGGAGGGTGAGTGGATCAGCACT**TTCCGCT**CATAACTGACTTCACCCATGGCTAG |
| mNZ38_13 | GTCCAAGCTTCTGGAGGGTGAGTT**AGCGGTA**ACGCTGCCTCGATAACTGACTTCACCCATGGCTAG |
| mNZ38_14 | GTCCAAGCTTCTGGAGGGTG**AGTCGCC**TAGCTCTCGAGTGTCATAACTGACTTCACCCATGGCTAG |
| mNZ38_15 | GTCCAAGCTTCTGGAGGGTGAGTAC**TGACGCT**ATGCCCGCCAATAACTGACTTCACCCATGGCTAG |
| mNZ38_16 | GTCCAAGCTTCTGGAGGGTGAGTAAACCCAGCA**TCCCGCG**CCATAACTGACTTCACCCATGGCTAG |
| mNZ38_17 | GTCCAAGCTTCTGGAGGGTGAGTCTGCTCCCCTAC**ACCCGCC**ATAACTGACTTCACCCATGGCTAG |
| mNZ38_18 | GTCCAAGCTTCTGGAGGGTGAGTTCCCT**CACCGCG**AGGTGGTATAACTGACTTCACCCATGGCTAG |
| mNZ38_19 | GTCCAAGCTTCTGGAGGGTGAGTAACCAG**GCACGCC**GCGTTGATAACTGACTTCACCCATGGCTAG |
| mNZ38_20 | GTCCAAGCTTCTGGAGGGTGAGTTGGCA**CAACGCC**ACCTGCTATAACTGACTTCACCCATGGCTAG |
| mNZ38_21 | GTCCAAGCTTCTGGAGGGTGAGTAGC**ATCCGCC**CATTTCCTAATAACTGACTTCACCCATGGCTAG |
| mNZ38_22 | GTCCAAGCTTCTGGAGGGTGAGTACTGGGCCCAAG**ACCCGCC**ATAACTGACTTCACCCATGGCTAG |
| mNZ38_23 | GTCCAAGCTTCTGGAGGGTGAGTGGT**AGCGTAT**GGCAACAGTATAACTGACTCCACCCATGGCTAG |
| mNZ38_24 | GTCCAAGCTTCTGGAGGGTGAG**TATCGCT**TTAAAC**ACTCGCT**ATAACTGACTTCCCCCATGGCTAG |
| mNZ38_25 | GTCCAAGCTTCTGGAGGGTGAGTAGCCTATC**ACCCGCA**GCCAATAACTGACTTCACCCATGGCTAG |
| mNZ38_26 | GTCCAAGCTTCTGGAGGGTGAGTC**ATACGCC**CGACGTCAGGAATAACTGACTTCACCCATGGCTAG |
| mNZ38_27 | GTCCAAGCTTCTGGAGGGTGAGTCGGCA**GGCGGAT**TCCCATTATAACTGACTTCACCCATGGCTAG |
| mNZ38_28 | GTCCAAGCTTCTGGAGGGKGAGTTG**TGCCGCA**CTATGTCTCCATAACTGACTTCACCCATGGCTAG |
| mNZ38_29 | GTCCAAGCTTCTGGAGGGTGAGTACTAGT**AGCCGCT**GCCGTTATAACTGACTTCACCCATGGCTAG |
| mNZ38_30 | GTCCAAGCTTCTGGAGGGTGAGTCCAGTAAGGTT**GGCGGCA**CATAACTGACTTCACCCATGGCTAG |
| mNZ38_31 | GTCCAAGCTTCTGGAGGGTGAGTATA**AGCCGCG**TTCCCTGCAATAACTGACTTCACCCATGGCTAG |
| mNZ38_32 | GTCCAAGCTTCTGGAGGGTGAGTATCTAGAACTAAG**GACCGCA**TAACTGACTTCACCCATGGCTAG |
| mNZ38_33 | GTCCAAGCTTCTGGAGGGTGAGTAGC**CATCGCA**ACGGTAAGCATAACTGACTTCACCCATGGCTAG |
| mNZ38_34 | GTCCAAGCTTCCGGAGGGTGAGTCACCAC**CTTCGCT**CGCCACATAACTGACTTCACCCATGGCTAG |
| mNZ38_35 | GTCCAAGCTTCTGGAGGGTGAGTACCAC**AGCGCGT**TGTGAGTATAACTGACTTCACCCATGGCTAG |
| mNZ38_36 | GTCCAAGCTTCTGGAGGGTGAGTAAACG**TTCCGCC**CGTGACCATAACTGACTTCACCCATGGCTAG |
| mNZ38_37 | GTCCAAGCTTCTGGAGGGTGAGTCCCCTA**CTCCGCT**AGTGGGATAACTGACTTCACCCATGGCTAG |
| mNZ38_38 | GTCCAAGCTTCTGGAGGGTGAGTCGATGCAACGG**GGCGGCC**CATAACTGACTTCACCCATGGCTAG |
| mNZ38_39 | GTCCAAGCTTCTGGAGGGTGAGTGTAA**TACCGCC**AACTCGATATAACTGACTTCACCCATGGCTAG |
| mNZ38_40 | GTCCAAGCTTCTGGAGGGTGAGTTGCCCATACA**CACCGCA**CTATAACTGACTTCACCCATGGCTAG |
| mNZ38_41 | GTCCAAGCTTCTGGAGGGTGAGT**TGACGCG**TACGGAATCCACATAACTGACTTCACCCATGGCTAG |
| mNZ38_42 | GTCCAAGCTTCTGGAGGGTGAGTGGCGTCAACC**TACCGCA**CCATAACTGACTTCACCCATGGCTAG |
| mNZ38_43 | GTCCAAGCTTCTGGAGGGTGAGT**GTACGCC**AATGACGTTGGCATAACTGACTTCACCCATGGCTAG |
| mNZ38_44 | GTCCAAGCTTCTGGAGGGTGAGTCGCATA**TCCCGCC**GCTTGGATAACTGACTTCACCCATGGCTAG |
| mNZ38_45 | GTCCAAGCTTCTGGAGGGTGAGTAGCCCGTAGCCT**TAGCGCC**ATAACTGACTTCACCCATGGCTAG |
| mNZ38_46 | GTCCAAGCTTCTGGAGGGTGAG**TGCGCCA**CCAACATTCTGTTATAACTGACTTCACCCATGGCTAG |
| mNZ38_47 | GTCCAAGCTTCTGGAGGGTGAGTTGCAGAAGAGAA**TAGCGCT**ATAACTGACTTCACCCATGGCTAG |
| mNZ38_48 | GTCCAAGCTTCTGGAGGGTGAGTGCCTTTACCCTG**TTACGCC**ATAACTGACTTCACCCATGGCTAG |

*Primer flanks are underlined in each sequence while the consensus sites are in bold.

**Table S2.** Oligonucleotide sequences utilized for NMR and EMSA

| | |
|---|---|
| mTMPRSS2_F | 5′-$^{1}$CACTGGGACAC$^{m}$CGCCTCCTGAG$^{22}$-3′ |
| mTMPRSS2_R | 5′-$^{23}$CTCAGGAGG$^{m}$CGGTGTCCCAGTG$^{44}$-3′ |
| TMPRSS2_F | 5′-$^{1}$CACTGGGACACCGCCTCCTGAG$^{22}$-3′ |
| TMPRSS2_R | 5′-$^{22}$CTCAGGAGGCGGTGTCCCAGTG$^{44}$-3′ |
| E-box_F (1) | 5′-$^{1}$GTTCAGAGGCAGGTGCCTGTGA$^{22}$-3′ |
| E-box_R | 5′-$^{23}$TCACAGGCACCTGCCTCTGAAC$^{44}$-3′ |
| mCZ38BS_21mer_F (2, 3) | 5′-$^{1}$GCACTCAT$^{m}$CGG$^{m}$CGCAGATCAG$^{21}$-3′ |
| mCZ38BS_21mer_R | 5′-$^{22}$CTGATCTG$^{m}$CGC$^{m}$CGATGAGTGC$^{42}$-3′ |

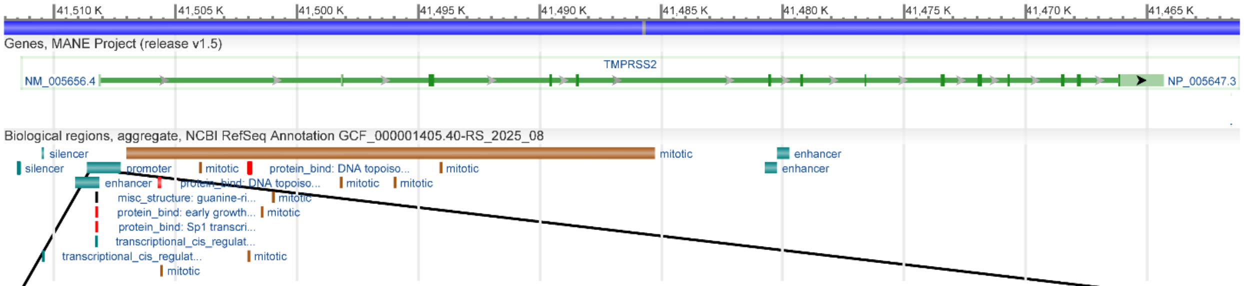


>NC_000021.9:41507255-41508648 Homo sapiens chromosome 21, GRCh38.p14 Primary Assembly

GCCGGGCAGGACAGGATGAGGTGGACCGAAGCGCCCAGGTGCCCCGCCTTCCCCCCAACCCGGCACGGAG
CGCAAGCGGAGGAGCCAGACCGGCTCCTCCCGGGCGGGTTCCTGCCGGTGCCGGGTGTCCAGGAGGCCCC
GGCCCTGATCGCCGCCAGCGGGTCGCCGCAACGGGGCCTGAGTCGCGAGCAGTCCCTCCCCGAGACTGAC
TGGCCGTTCGCACCTCGCCCGGGGGCCCTCCAGCAGCCTGGCCCAGCGGTCCCCGCCGCGCCGGCCGTAT
CGTTCCCGGCATCTCAGCGAGTTTCCAGTTTCTCCCTACCTCAGTAGGGAACCAAGAAGCAGAAGGAGTT
CAGGTAACACTGCGGGACTGCAGAGATTTCCCCCAAACAAAGAAGCCA**ACCGAC**AGCACTCCCAGTCCTC
CTCCCCAAAGAGAAAAGGCGCACCGGTGCTCCCAGGCGGGGGCCGTGGAGGGCAGGCGGACTAGGAGCCA
GCTTTGGGGACCCCGGGGGACTCTCTTCCACCAACTGGCTAGGGACTCGCAGGACCACCTGCCCCAACCT
CGCGATGCCAAGCCAACAGGGGCGGCGAGGGCTCTCGGCCGTGCGCAAGGGGTCCCAGGCGCCCAGCACT
CTCCCAGCACCCCGGGAGGCGCCCTGCCCGGCTGGCCCCAGCGCTCGACCCTCGGGCGCACTCACCTGCC
GCGCCGCGCTCCTCACACCCGCTTTCACCTCCGGGCGGGGCAGGGGGCATCGGCGGGTCCCAGGCGCCCA
GGTTCCCCTCCCCAGCCCGGACCCCGAGCCGGGACCCTGGTACCGGCGCCGCTCACCTGCCGCGCTCCAG
GCGGCGCTCCCCGCCCCTCGCCCTCCGCCTCCGCCTCCGCCTCCTGCTTAGCTCGCGCCTACTCGGCCCG
GCCCGCCCTGGCTCTCGCTTTAATCTCAGGA**GGCGGTG**TCCCAGGGTCCAGGGCGACACCCCCAAGCGGG
TCGCGGGTCACCCTGGGCGGGCGGGGGCGGGGGCGGCGGGAGGAGGCGTTTCCCAAGGGAAGTCCGGGAC
GGGTTCCCTGCCTGGCTCAACCCCGGCTGCGGAGGGACCCATAAGGGACTCTGGGGCGCTTCCCGGCCC
GGCCACAGCGGCCAACCTGCCCCTCTCCGGTCCGCGGCGCAGCGCTGCCCGAGGGCTCGAGTTTGGGTTA
AGGAAGGGGATTCTGGGGAGCGCTCCTGGAGTCCTGGGTGAGCCCCCGGAGCTGCTACCTCACGAGCTCC
TGTAGGCGGCCGGCCCCGTCTTGGCCTCCGGGGAACCCACTGTAGCCTCTGGGCAAGGACCCAGGGTAGG
CCTTGGGGACTGCTGGCCCAAGCACGAGTGGCGGGGACACAGCGCCGGACGCCCAAGACCCTCC

**Figure S1.** Gene annotation for the human TMPRSS2 gene from the current GRCh38.p14 primary assembly (Ch 21; 41,508,158 – 41,464,305). The TMPRSS2 promoter FASTA sequence (41,507,255 – 41,508,648) (4) is depicted below, denoting the locations of the ZBTB38 N- (*blue highlight*) and C-terminal (*green highlight*) ZF domain consensus sequences. The underlined region was used to design the mTMPRSS2 DNA target utilized in this study (Table S2). The top figure was generated using the NCBI Genome Data Viewer (reference sequence NC_000021.9) (5).

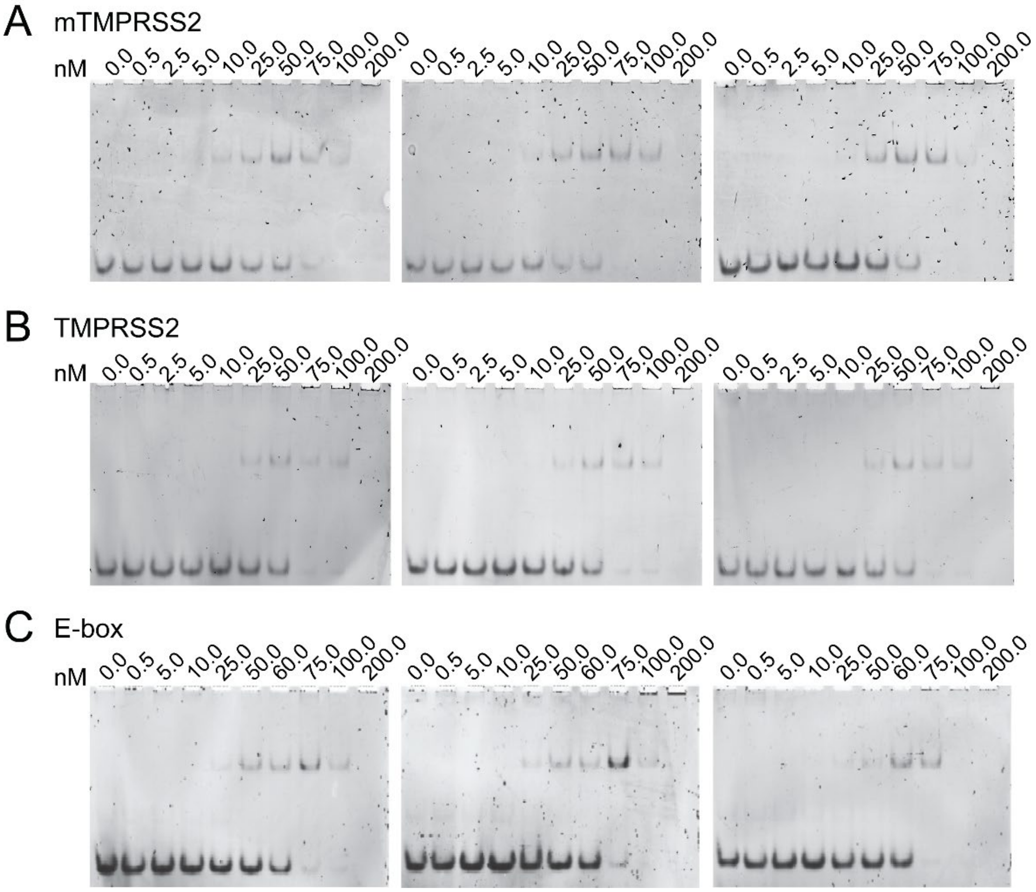


**Figure S2.** EMSA gel triplicates for ZBTB38 ZFs 1-5 binding to *A*, mTMPRSS2 (mCpG); *B*, TMPRSS2 (CpG); and *C*, E-box (TpG). These gels were used to generate binding isotherms and determine $K_d$ values associated with Fig. 2C.

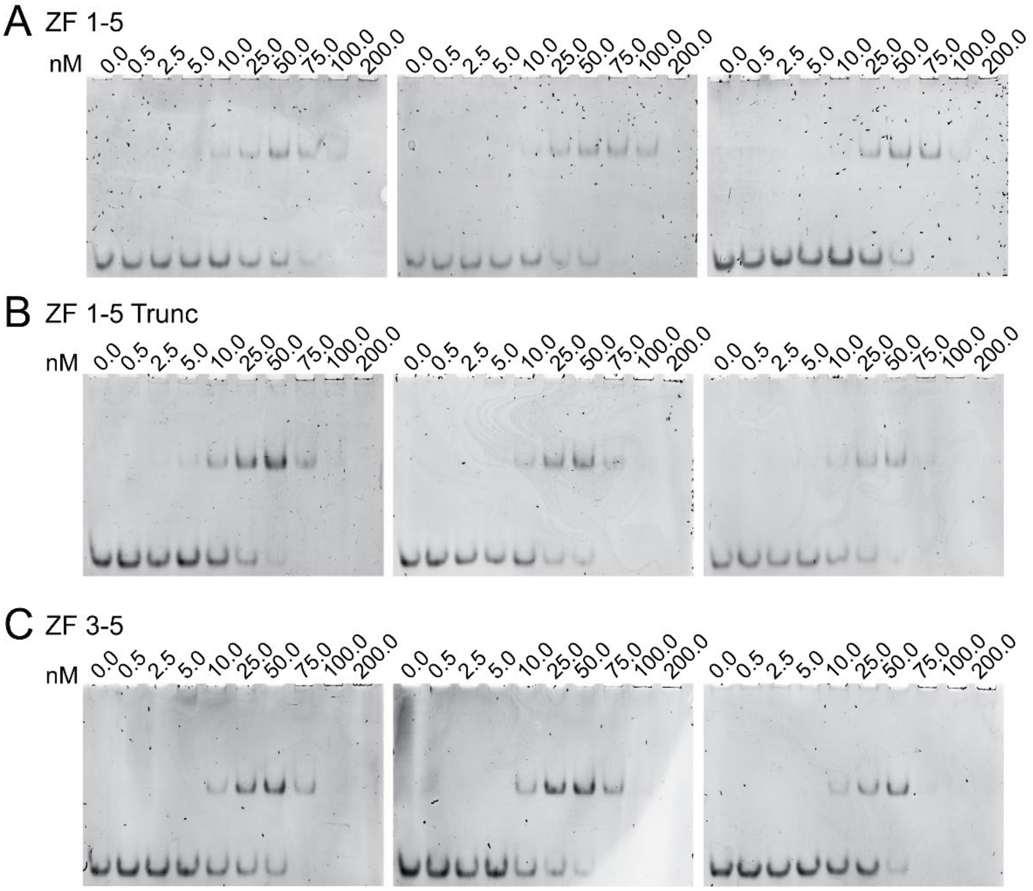


**Figure S3.** EMSA gel triplicates for ZBTB38 *A,* ZF 1-5; *B*, ZF 1-5 Trunc; and *C*, ZF 3-5 binding interactions with mTMPRSS2. The gels in *A* are reproduced from Fig. S2A for comparison purposes. These gels were used to generate binding isotherms and determine $K_d$ values associated with Fig. 3D.

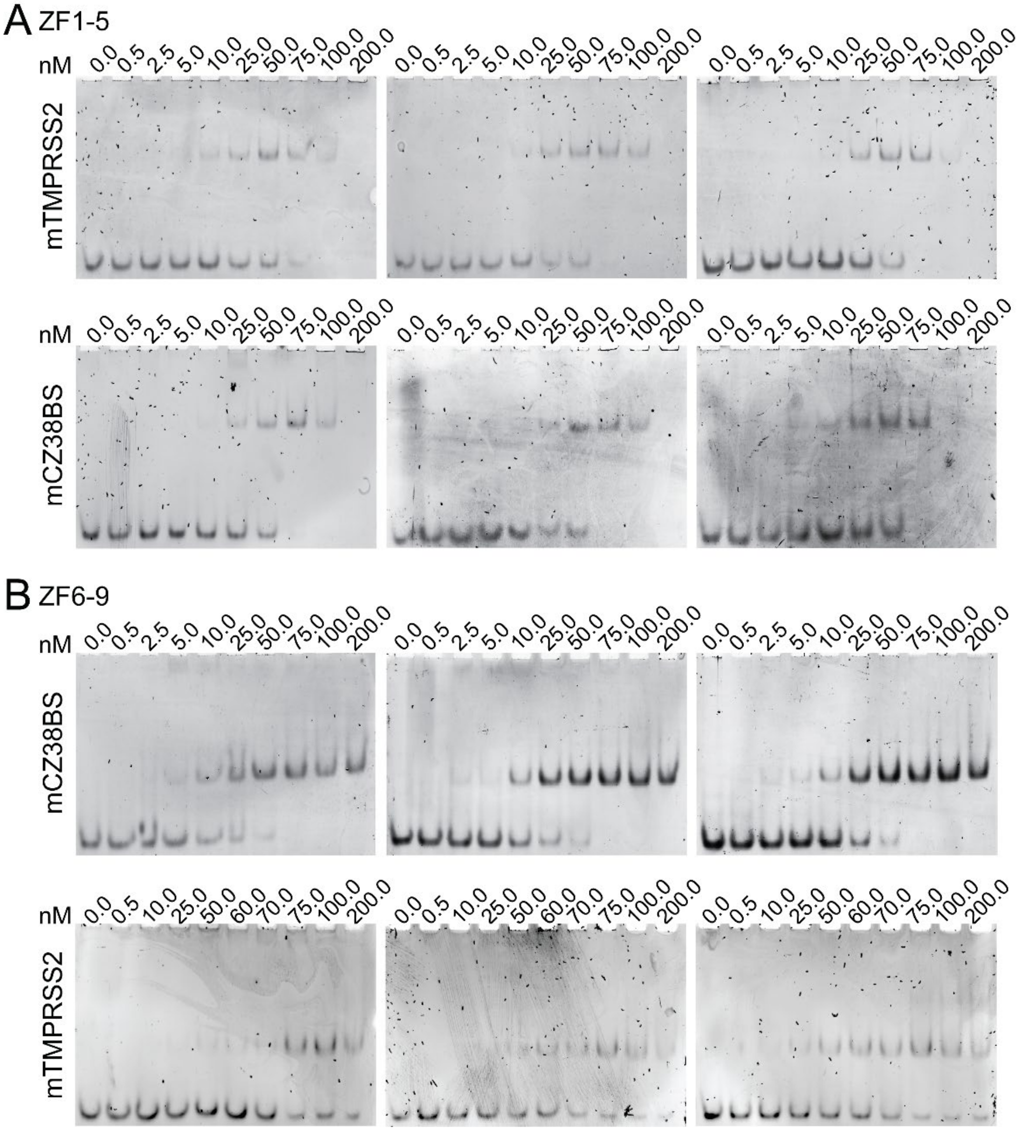


**Figure S4.** EMSA gel triplicates for *A*, ZBTB38 ZF 1-5 binding to mTMPRSS2 (*top*) or the ZBTB38 C-term ZF consensus sequence (mCZ38BS (3); *bottom*) and *B*, ZBTB38 ZFs 6-9 binding to mCZ38BS (*top*) or mTMPRSS2 (*bottom*). The ZF 1-5:TMPRSS2 gels in *A* are reproduced from Fig. S2A for comparison purposes. These gels were used to generate binding isotherms and determine $K_d$ values associated with Fig. 4.
